# Distinct spatial and non-spatial response properties of excitatory narrow-spike and burst-firing neurons in the marmoset auditory cortex

**DOI:** 10.64898/2026.02.10.705046

**Authors:** Chenggang Chen, Evan Remington, Xiaoqin Wang

**Affiliations:** Laboratory of Auditory Neurophysiology, Department of Biomedical Engineering, Johns Hopkins University School of Medicine, Baltimore, Maryland 21205, USA

## Abstract

Cell types are the basic units of the cerebral cortex. Broad and narrow spike waveforms (BW and NW) are the primary criteria for classifying putative excitatory and inhibitory cortical neurons, particularly in non-human primates where genetic accessibility is limited. We have classified cell types in the auditory cortex of marmosets using spike waveforms and firing patterns and identified a new type of neuron: the NW-burst neuron. NW-burst neurons are excitatory, as they drive but do not suppress the firing of connected neurons. Furthermore, NW-burst neurons displayed shorter response latencies, lower response variability, smaller receptive fields, stronger correlations between spatial and non-spatial selectivity, and higher decoding accuracy than other neurons. Together, these findings suggest that cortical NW-burst neurons in non-human primates represent a distinct excitatory cell type.

**Significance Statement:** Classifying cell types in non-human primates is challenging due to limited genetic tools. Traditionally, neurons with narrow spikes are assumed to be inhibitory. Challenging this view, we analyzed 1,816 high-quality single units in the marmoset auditory cortex and discovered that narrow-spike burst-firing neurons were excitatory. These neurons exhibited shorter response latencies, smaller receptive fields, and higher decoding accuracy. Crucially, they showed a positive correlation between "where" and "what" selectivity, defying the standard trade-off between spatial and non-spatial response properties. These findings reveal a specific excitatory cell type in the auditory cortex of a non-human primate species that integrates sound information with high temporal fidelity and feature selectivity.

## Introduction

Cell types exhibit diverse properties in many modalities—molecular, morphological, and electrophysiological (Zeng, 2022). Cortical computations, such as attention, decision making, and sensory processing, depend on the interactions between excitatory and inhibitory neurons (Harris and Mrsic-Flogel, 2013). Before we can correlate cortical computations with cell types, we first need to classify them. For longitudinal electrophysiological recordings in primates, the primary marker is the spike waveform: putative excitatory and inhibitory neurons have broad and narrow spike waveforms (BW and NW), respectively. Numerous studies show that BW and NW neurons exhibit distinct functions in the macaque (Diester and Nieder, 2008; Vinck et al., 2013; Dasilva et al., 2019; Milton et al., 2020; Roussy et al., 2021) and marmoset (Gilliland et al., 2024; More et al., 2024) cortex. In macaques, NW neurons show stronger response reliability during attention in V4 (Mitchell et al., 2007) and are more selective for auditory categories in auditory cortex (Tsunada et al., 2012). In marmosets, NW neurons show lower stimulus selectivity in auditory cortex (Gao and Wang, 2018) and earlier stimulus-related activity in parietal cortex (Selvanayagam et al., 2024).

In addition to waveform, firing pattern can also be used to classify cell types. One example is bursting neurons, which fire spikes rapidly in a short time period (Bair et al., 1994; Livingstone et al., 1996; Friedman-Hill et al., 2000; Katai et al., 2010; Womelsdorf et al., 2014). Firing bursts have many advantages over firing isolated spikes (see review Lisman, 1997; Zeldenrust et al., 2018). They reliably represent and transmit a stimulus (Martinez-Conde et al., 2002; Naud and Sprekeler, 2018), and can modify brain state (Li et al., 2009), induce synaptic plasticity (Sjöström et al., 2001), and coordinate learning in deep networks (Payeur et al., 2021). Bursting neurons typically have a BW (Ardid et al., 2015; Trainito et al., 2019). There is a unique type of high-frequency (350 to 700 Hz, Nowak et al., 2003) bursting neuron that surprisingly has a NW (i.e., NW-burst). NW-burst neurons are found in macaque and capuchin V1 (Onorato et al., 2020), macaque V4 (Mitchell et al., 2007; Anderson et al., 2011), macaque prefrontal (Hussar and Pasternak, 2009) and motor (Chen and Fetz, 2005) cortex, and marmoset auditory cortex (Liu and Wang, 2022), but not in mouse V1 (Onorato et al., 2020).

Morphological evidence suggests that NW-burst–like “chattering” neurons in cat V1 are excitatory (Gray and McCormick, 1996; Nowak et al., 2003; Cardin et al., 2005). However, Katai et al. (2010) report both excitatory and inhibitory high-frequency (>200 Hz) bursting neurons in macaque frontal cortex. Rodent studies show that cortical NW inhibitory parvalbumin (PV) neurons fire bursts of spikes (70 to 260 Hz, Puig et al., 2008; Deleuze et al., 2019; Lee et al., 2021). Basket (fast spiking), chandelier, and multipolar bursting neurons (Baltow et al., 2003) are the three major types of PV neurons (see review Tremblay et al., 2016). Burst firing is also found in other types of inhibitory neurons, including somatostatin (SST; low-threshold spiking or Martinotti, Kawaguchi and Kubota, 1996; Goldberg et al., 2004; Ma et al., 2006), vasoactive intestinal peptide (VIP; Lee et al., 2010; Prönneke et al., 2015), and calretinin (Caputi et al., 2009) neurons. Therefore, it remains unclear whether the unique NW-burst neurons in primates (or larger mammals) are excitatory or inhibitory.

In this study, we used a combination of spike waveform and firing pattern analyses to classify three cell types: NW-burst, NW-nonburst, and BW (burst and nonburst) based on over 1800 single units recorded from awake marmoset auditory cortex. Using paired-neuron recordings, we validated that NW-burst neurons are excitatory. We further corroborated this by comparing the auditory spatial, spectral, and intensity receptive fields between connected pairs. Finally, we demonstrated distinct spatial and non-spatial tuning properties among the three types of neurons.

## Materials and Methods

### Electrophysiological recordings in marmosets

All experimental procedures were approved by the Institutional Animal Care and Use Committee of Johns Hopkins University and followed National Institutes of Health guidelines. We recorded well-isolated single-unit activity in the auditory cortex of four marmoset monkeys (Callithrix jacchus). A total of 1,816 units were recorded: 326 from M3T, 732 from M71V, 455 from M9X, and 303 from M6X. We previously published spatial receptive fields (SRFs) from 649 of these units, recorded in M3T (164), M71V (311), and M9X (174) under passive listening conditions (Remington and Wang, 2019). From this dataset, 209 single units recorded in M71V and M9X during a sound localization task were also reported (Chen et al., 2025a). Recordings were obtained from primary auditory cortex (A1), the rostral area (R), the rostrotemporal area (RT), and caudal medial/lateral belt areas (CM/CL), with boundaries identified using frequency reversals. In this study, we did not analyze differences across cortical areas.

Single-neuron activity was recorded with a single-electrode preparation in the left auditory cortex of three awake marmosets and right auditory cortex of one marmoset (M6X). Animals were trained to sit in a custom-designed primate chair. After acclimation, two stainless steel headposts were surgically implanted under sterile conditions to stabilize head orientation during recordings. To access the auditory cortex, small craniotomies (∼1 mm in diameter) were made over the superior temporal gyrus. Tungsten electrodes (2–5 MΩ impedance; A-M Systems) were advanced with a hydraulic microdrive (Trent-Wells). Single-unit activity was sorted online using a template-based spike-sorting system (MSD; Alpha Omega Engineering) and analyzed offline with custom MATLAB code (MathWorks). Throughout the sessions, an infrared video camera was used to monitor the state of the animals.

Stimuli were generated in MATLAB at a sampling rate of 97.7 kHz using custom software. Digital signals were converted to analog (RX6, 2-channel D/A; Tucker-Davis Technologies, TDT), attenuated (PA5 ×2; TDT), power-amplified (Crown Audio ×2), and delivered through a selected channel of a power multiplexer (PM2R ×2, 16 channels; TDT). Amplifier gains were adjusted such that the intensity of a 4 kHz tone presented from the front speaker was 95 dB SPL at 0 dB attenuation, measured with a 0.5-inch free-field microphone (Brüel & Kjær model 4191) positioned at the animal’s head location. Loudspeaker responses were measured using Golay codes and exhibited relatively flat frequency response curves (±3–7 dB) and minimal spectral variation across speakers (<7 dB relative to mean) across the frequency range of the stimuli used (Chen et al., 2025b).

When possible, neurons were characterized for frequency, intensity, and spatial tuning. Characterization was performed by recording responses to sets of stimuli, typically 200 ms in duration, presented in pseudorandom order. Each stimulus was delivered 5–10 times. For frequency tuning, stimuli consisted of pure tones and bandpass-filtered Gaussian noise. The frequency axis was sampled in 0.1-octave steps, typically spanning a four-octave range (2–32 kHz). Firing rates were calculated within a window beginning 15 ms after stimulus onset and ending 20 ms after stimulus offset. All stimuli used to measure spatial receptive fields were either bandpass-filtered or constructed to have energy between 2 and 32 kHz. Vocalization stimuli consisted of ten types of natural and time-reversed marmoset calls, as in our previous studies (Liu and Wang, 2022).

For the attentive listening condition, we implemented a Go/No-Go discrimination task (Chen et al., 2023). In this task, animals were trained to respond with a lick at the feeding tube to target sounds in order to receive a food reward, while withholding responses when no target was presented. Each trial consisted of two phases: a variable-length intertrial interval and a fixed-length response interval. During the intertrial interval, sounds were presented only from background locations. The length of the intertrial interval was randomized between approximately three and ten stimuli. The response interval always included four alternations between target and background locations. If an animal responded during the intertrial interval, the trial was aborted, followed by a time-out (and occasionally a mild air puff to the base of the tail), after which the intertrial interval restarted. At the end of the intertrial interval, target sounds alternated with background sounds during the response interval. Trials ended either when the response interval expired or when a lick was detected during the response interval. Correct responses were rewarded with approximately 0.1–0.2 ml of food. If no lick was detected, the next trial began immediately. To measure false alarms, one third of trials were sham trials in which stimulus location did not change. In these trials, licks were not rewarded.

### Cell type classification

We collected all the single units using one tungsten electrode. We recorded high-quality single units by manually matching their spike waveform with a template. The online template matching method allows us to continually collect the same waveform from a single unit. Therefore, there is no noisy spike, and spike sorting is unneeded. One limitation of this method is the limited recording time since some neurons will drift away or be “killed” by the electrode when it is too close. Therefore, we can only collect a much smaller number of spikes for each unit. This will make the calculation of ACG and CCG unreliable if number of spikes are limited. To make the calculation robust, we removed those units that have less than five spikes at the peak of ACG and got 1098 out of 1816 units. For the ISI histogram shown in Supplementary Figure 1b, the X axis (1 to 80 ms) was 10 to the power of 0:0.05:log10(80), same as Nowak et al., 2003. Therefore, the size of each time bin was not linear. This was different from the linear 0.2 ms time bin in the ACG histogram (X-axis: 0.2 to 100 ms). The ACG decay measures how strongly the auto-correlogram (ACG) falls off after its peak. First, identify the ACG peak at time p (in milliseconds). Second, compute the average ACG value in a small window around the peak, from (p−1 ms to p+3 ms). Call this the local peak average. Third, compute the average ACG value over a later baseline window, from 30 to 100 ms. Call this the baseline average. Finally, the ACG decay percentage is then defined as the difference between the local peak average and the baseline average, divided by their sum (see Fig. 1c).

**Figure 1.**
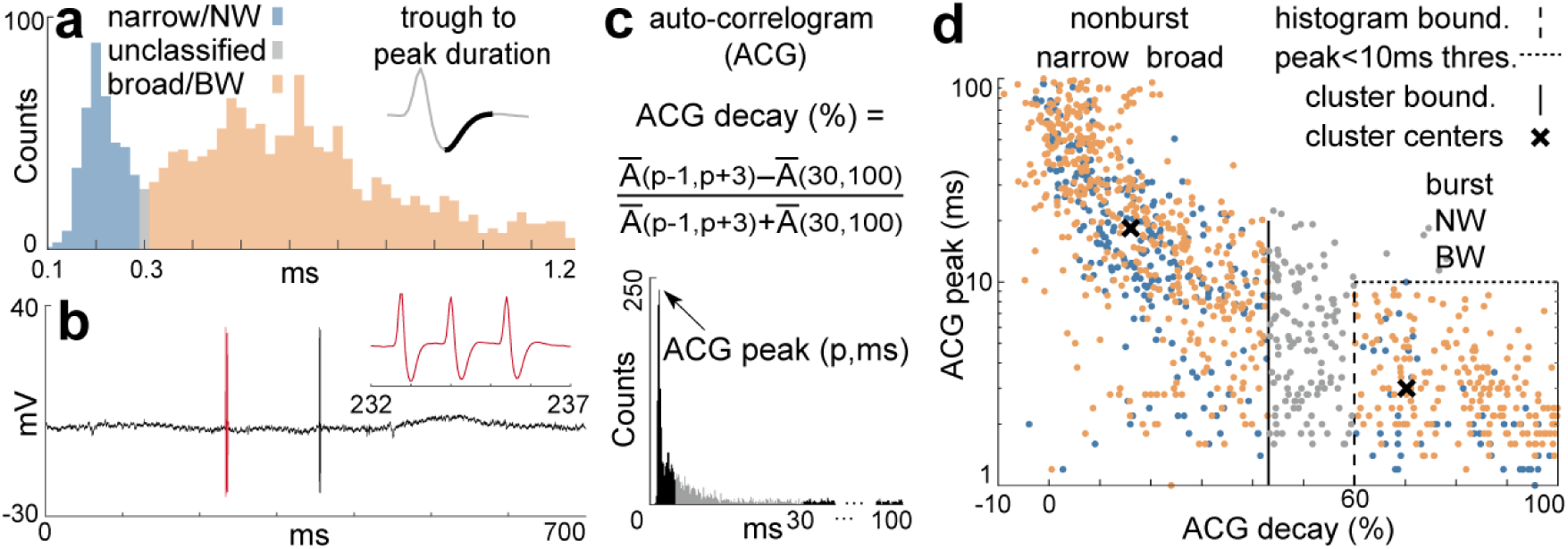
Classifying cell types using spike waveforms and firing patterns. (a) Distribution of trough-to-peak duration in marmosets (bin width = 0.02 ms). The numbers of NW, unclassified, and BW units are 385, 26, and 1405 (1816 in total). (b) Raw spike waveforms of a single unit (M71V-4283) in response to a sound stimulus. Inset: three spikes occurring within 5 ms of the highlighted time period. (c) Calculation of ACG peak and decay from the same unit. (d) Scatter plot of ACG peak versus decay in marmosets. A total of 1099 units were included in the analysis. The numbers of NW-nonburst, BW-nonburst, NW-burst, and BW-burst units were 250, 469, 54, and 199, respectively. The cluster boundary is 43%, with cluster centers at (nonburst: 16.4%, 18.4 ms) and (burst: 70.3%, 3 ms).

### Cell type validation

Our online spike waveform template matching software allows us to record at most three units simultaneously. We identify monosynaptic excitatory or inhibitory interactions between simultaneously recorded neuron pairs using the spike cross-correlogram (CCG). We used the same “xcorr” function in MATLAB to compute both ACG and CCG. ACG only needs one input, i.e., the spike train of one neuron. CCG requires two spike trains from paired neurons, along with the maximum delay (-50 to 50 ms). We identified any significant connection using the spike jitter method. We chose a time bin of 1 ms and a jitter range of -5 to 5 ms. We repeated the calculation 500 times. CCG will be different because the spike train of post-synaptic target neuron was jittered. We computed the 99% (2.576 standard deviation, std), 99.9% (3.291 std), and 99.99% (3.891 std) confidence intervals. We also computed the global threshold, which was the maximum and minimum values of 99.99% confidence intervals. If the original, without jittering CCG, was higher or lower than the global threshold, then the pre-synaptic neuron will be validated as excitatory or inhibitory, respectively. In other words, a significant peak in the CCG indicates an excitatory drive connection, while a significant valley indicates an inhibitory suppress connection.

### Response reliability, spatial receptive fields, and sound decoding

Response reliability: We used the MATLAB code provided by Mark Churchland’s lab (Variable toolbox, https://churchland.zuckermaninstitute.columbia.edu/content/code). The authors provided a detailed tutorial on how to use their code. We did not modify their code or the parameters. The only user-adjustable parameter is “boxWidth,” that used to slide (or smooth) the curves. Longer time windows, like 100 ms, make real effects larger, and reduce the impacts of artifacts related to nonIZPoisson spiking. However, it will reduce the time resolution. We used the default 50 ms. It also requires the alignment time, which is the stimulus onset time. We used 200 ms (pre-stimulus time). We computed the Fano factors for three cell types separately using the provided function “VarVsMean”. This function essentially regresses the spike-count variance versus the spike-count mean and reports the slope (i.e., Fano Factor). The input data are the spike raster of each trial with a time resolution of 1 ms. For example, a stimulus with 10 repetitions and 700 ms ISI will generate a 10x700 logical array. All 24 stimuli from all single units will be combined, which results in 792, 2064, and 9600 logical arrays for NW-burst, NW-nonburst, and BW neurons.

Characterization of spatial receptive fields: To partially alleviate biases in calculated tuning properties that could be introduced by uneven spacing of speakers, spatial receptive fields were generated by interpolating responses to the 24-speaker array into a 5° × 5° (2592 zone) vertical pole grid according to the spherical distance-weighted mean of the two nearest speaker locations. For each interpolated grid location, the contribution (weight) of a nearby speaker is determined by the inverse square of the spherical distance between the speaker and the grid center. Each weight is then normalized by dividing it by the sum of the inverse squared distances of the two closest speakers. This ensures that speakers closer to the grid location have proportionally more influence than farther ones. The tuning area describes the proportion of the spatial grid where the neuron responds above a threshold. For each grid zone, if the interpolated firing rate is higher than the threshold (defined as half of the maximum firing rate), the area of that grid zone is added. The sum of these responsive areas is then divided by the total area of the grid. A neuron that responds everywhere has a tuning area of 1, while one that responds only to a single location has a very small tuning area (around 0.045).

Sound location decoding: For a single neuron, the probability p given that a response r was evoked by an ILD stimulus s was decided by Bayes’ rule as:

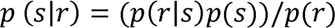

Since the probability of the presented stimuli *p*(*s*) was 1/24, and *p*(*r*) does not affect the selection of the stimulus that maximizes the posterior probability. Thus, *p* (*s*|*r*) was proportional to *p*(*r*|*s*), i.e., one of the 24 stimuli that have the highest probability to evoke this response. The stimulus that has the largest probability was chosen and will be compared with the real stimulus. The chance level was 4.17%. At each repetition of a fixed number of neurons, if six out of 24 stimuli have been successfully predicted, the correct percentage should be 25%. We used the population pattern decoder same as the one used in our previous paper (Chen and Song, 2024):

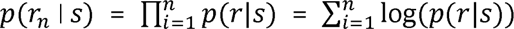

Here, we used the maximum likelihood estimation (MLE) (Miller and Recanzone, 2009) decoding method, which assumes Poisson distributions of neural responses. We run the decoder 100 times for each cell type separately. We showed the mean and standard deviation among 100 repetitions in Figure 6b. The mean values were smoothed using the “smooth” function in MATLAB (span = 0.15).

## Results

This study is based on 1,816 well-isolated single units recorded from the auditory cortex of four marmosets that were either passively listening or engaged in a sound localization task (Fig. 1a). Using a single tungsten electrode and online template matching, we recorded 1 to 3 units simultaneously. While most recordings yielded single units, we obtained 336 pairs and, more rarely, 44 triplets. Spatial tuning and dynamic properties of a subset of these data (649 units) were reported in our previous studies (Remington and Wang, 2019; Chen et al., 2025a). Under the passive listening condition, we measured auditory spatial receptive fields (SRFs) in all four marmosets (Remington and Wang, 2019). Broadband noises were delivered from 24 speaker locations that covered the full field around the marmoset’s head. Other stimuli, including pure tones and vocalizations were also presented. Under the active listening condition, two of the four marmosets heard a continuous sound from one target speaker location and were trained to discriminate sound location changes among four target speaker locations (Chen et al., 2025a).

### Classification of cell types based on spike waveforms and firing patterns

The clearest and most widely used criterion for classifying cell types is the width of the spike waveform: neurons with a narrow waveform (NW) are generally considered inhibitory, whereas neurons with a broad waveform (BW) are considered excitatory. We used the trough-to-peak duration (Fig. 1a, inset) as the metric to separate NW and BW neurons. In marmosets, the distribution of trough-to-peak duration was bimodal (Hartigan’s dip test, p < 0.001, amplitude: 0.025), with a valley at 0.3 ms (gray bar, Fig. 1a). Using this boundary, we classified 1,816 neurons into 21.2% NW neurons (blue bars) and 77.4% BW neurons (orange bars).

Different cell types also exhibited distinct patterns of spiking. In addition to regular spiking in excitatory neurons and fast spiking in inhibitory neurons, another characteristic spiking pattern is bursting. Bursting neurons fire spikes very sparsely (Fig. 1b; Supplementary Fig. 1a), which distinguishes them from fast-spiking neurons. Bursting refers to intensive firing within a very short time window (Fig. 1b, inset), so the inter-spike-interval (ISI) histogram shows a sharp peak (1.7 ms, Supplementary Fig. 1b). In addition to ISI, the auto-correlogram (ACG) can also quantify differences in firing patterns, such as the strength of burstiness. The ISI histogram reflects the distribution of consecutive spike intervals, whereas the ACG reflects the histogram of all pairwise lags, not only consecutive ones. We used the ACG because its shape is a reliable marker for putative excitatory or inhibitory neurons (Barthó et al., 2004; Peterson et al., 2021) and can be combined with cross-correlograms to validate cell types (next section). For example, bursting neurons show a large peak at less than 10 ms followed by an exponential or sharp decay (Fig. 1c). In contrast, regular and fast-spiking neurons show an exponential rise (Supplementary Fig. 2, #8) or a sharp rise (Fig. 2b) from time 0, respectively. Because bursting neurons fire only a few spikes within a brief window, the ACG peak occurs at a short lag (1.68 ms), but the ACG decay is large (87.9%) (Fig. 1c).

**Figure 2.**
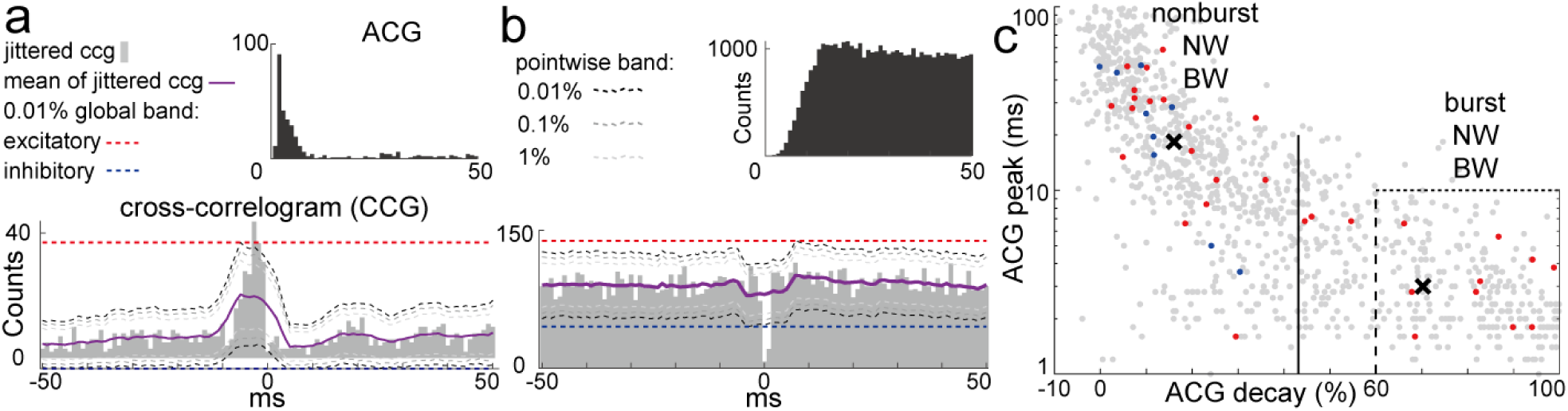
Validating the cell type classification using CCG. (a) ACG and its excitatory connection (CCG) of a putative excitatory neuron (M71V-0386; presynaptic and postsynaptic units were recorded from channel 5 and 6 of same tungsten electrode, respectively). (b) Same as (a), but for a putative inhibitory neuron with an inhibitory connection (M3T-2550; presynaptic and postsynaptic are channel 5 and 4, respectively). (c) Overlay of excitatory (red) and inhibitory (blue) neurons identified through CCG.

We used two metrics—the peak time in the ACG (ACG peak) and the ACG decay—to classify burst and nonburst neurons. Bursting neurons had an ACG peak of less than 10 ms and an ACG decay greater than 60%. The first criterion required a peak firing rate above 100 Hz, which largely excluded regular-spiking neurons. A cutoff of 10 ms has been used in many previous studies (de Kock Sakman, 2008; Onorato et al., 2020; Liu and Wang, 2022). The second criterion required a sparse firing rate, which largely excluded fast-spiking interneurons. A cutoff of 60% was determined from the histogram of ACG decay (Supplementary Fig. 1c). This threshold was conservative, as unsupervised K-means clustering (distance: squared Euclidean) separated burst and nonburst neurons at 43% (Supplementary Fig. 1d). We excluded all neurons with an ACG decay greater than 43% but less than 60% (gray dots in Fig. 1d).

With one criterion for NW/BW neurons and two criteria for burst/nonburst neurons, we classified neurons into four groups (Fig. 1d). NW-nonburst (blue) and BW-nonburst (yellow) neurons mainly occupied the top left corner, representing neurons with long intervals between pairwise spikes (y-axis) and dense firing (x-axis). NW-burst and BW-burst neurons occupied the bottom right corner, representing neurons with short intervals between pairwise spikes and sparse firing. Our main interest was to determine whether NW-burst neurons are inhibitory or excitatory, and to characterize their tuning properties. We therefore compared NW-burst neurons with the other NW type (NW-nonburst) and with BW neurons. The numbers (percentages) of NW-burst, NW-nonburst, and BW neurons were 54 (4.9%), 250 (23%), and 668 (61%), respectively.

### Validation of cell types based on functional connectivity

A major limitation of previous primate cell-type studies is the lack of validation. Here, we validated excitatory and inhibitory neurons using cross-correlograms (CCGs) between pairs of connected neurons recorded from marmoset auditory cortex (Fig. 2). The CCG measures the time delay between spikes of two neurons. If a postsynaptic target neuron consistently fires after a short delay from a presynaptic source neuron, then the source neuron is excitatory. Figure 2a shows an example CCG where a significantly higher number of spikes (above the red line) was observed within a 10 ms delay. In contrast, if a target neuron consistently suppresses firing after a source neuron spike, then the source neuron is inhibitory. Figure 2b shows an example CCG where a significantly lower number of spikes (below the blue line) was observed within a 10 ms delay.

To reach statistical significance, both source and target neurons must fire sufficient spikes, particularly for identifying inhibitory connections. Because excitatory neurons generally fire fewer spikes, connections from inhibitory to excitatory neurons are likely underestimated compared with inhibitory-to-inhibitory connections. Two observations support this interpretation. First, most target neurons were putative inhibitory. Supplementary Figure 2 shows ACGs and CCGs for all nine inhibitory-type connections across fifteen sessions. All source neurons displayed features of inhibitory neurons, i.e., sharp decay toward time 0 (except #8, which showed exponential decay). Interestingly, seven target neurons in thirteen sessions were also putative inhibitory. Second, a higher proportion of identified inhibitory neurons had a BW (Supplementary Fig. 1e, f). Among inhibitory neurons, NW parvalbumin (PV) neurons mainly inhibit excitatory neurons, a connection underestimated by CCG due to the low firing rates of excitatory neurons. In contrast, somatostatin (SST) and vasoactive intestinal peptide (VIP) neurons mainly have a BW and inhibit other inhibitory neurons. The unusually high percentage of identified inhibitory BW neurons may reflect this connection pattern.

We overlaid nine inhibitory (blue) and 31 excitatory (red) neurons on a scatter plot of ACG decay versus ACG peak (Fig. 2c). All inhibitory neurons were non-bursting, whereas 10 (32%) excitatory neurons were bursting (4.9% NW-burst and 18.1% BW-burst, 23% in total). Taken together, CCG data from marmoset auditory cortex supports the conclusion that NW-burst neurons are excitatory.

### Corroboration of the cell type validation by examining the tuning of paired neurons

One concern with using firing patterns to classify cell types is that brain state can modulate these patterns. For example, Anderson et al. (2013) reported that the burstiness of macaque V4 neurons is reduced by attention. To address this, we compared the firing patterns of individual neurons (ACGs) and the interactions between paired neurons (CCGs) in active (Fig. 3a) and passive (Fig. 3b) listening marmosets. We found that both ACGs and CCGs were qualitatively similar across conditions, though not quantitatively identical. Inhibition strength was larger under the behavioral condition. This difference may reflect brain state changes or the longer recording times (and thus more spikes) obtained during attentive sessions. Although the proportion of connected neurons may be underestimated in the passive condition, the sign of functional connectivity remained consistent across states. In addition to computing the lead–lag relationship of paired neurons, we also compared their sensory representations to corroborate the validation results. Specifically, we examined the auditory spatial, spectral, and sound level tuning of paired neurons. Depending on the sign of the CCG, the sound tunings were either complementary for inhibitory CCGs or overlapping for excitatory CCGs.

**Figure 3.**
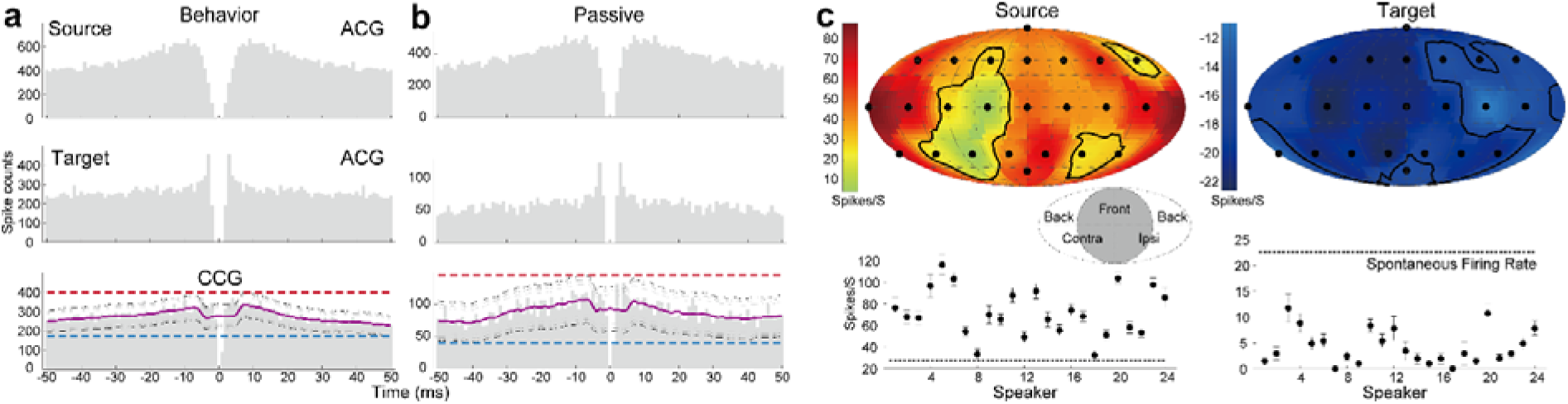
CCG is stable across brain states and consistent with sound tunings. (a) ACGs from source and target neurons (M71V-0566), and the CCG between them, during a spatial discrimination task. Note that the number of trials in the active condition was greater than in the passive condition. (b) Same neurons and analysis as in (a), but recorded while the animal was passively listening. (c) SRFs and firing rates of the source and target neurons shown in (a, b).

Figure 3c shows the spatial receptive fields (SRFs) and firing rates at each speaker location for a pair of neurons whose ACGs and CCGs are shown in Fig. 3a, b. The target neuron exhibited a similar spontaneous firing rate as the source neuron (Fig. 3c, bottom; Supplementary Fig. 3a). However, during sound stimulation, its responses were strongly suppressed by the source neuron, resulting in an SRF with negative firing rates across all locations (Fig. 3c, right). Notably, the target neuron remained selective for sound location (ANOVA, p < 0.001).

Figures 4a, b show spatial, spectral, and sound-level tuning for another pair in which the same source neuron inhibited the same target neuron. The source and target neurons were located in the contralateral and ipsilateral hemispheres of the SRF, respectively (Fig. 4a; Supplementary Fig. 3b). Although the two neurons had similar spontaneous firing rates (Fig. 4b, dashed lines), the target neuron (green) was strongly suppressed at frequencies above 8 kHz (Supplementary Fig. 3c), corresponding to the region where the source neuron (yellow) was most strongly activated. For sound levels below 60 dB attenuation (Fig. 4b, right; Supplementary Fig. 3d), the source and target neurons showed strong excitation and inhibition, respectively.

**Figure 4.**
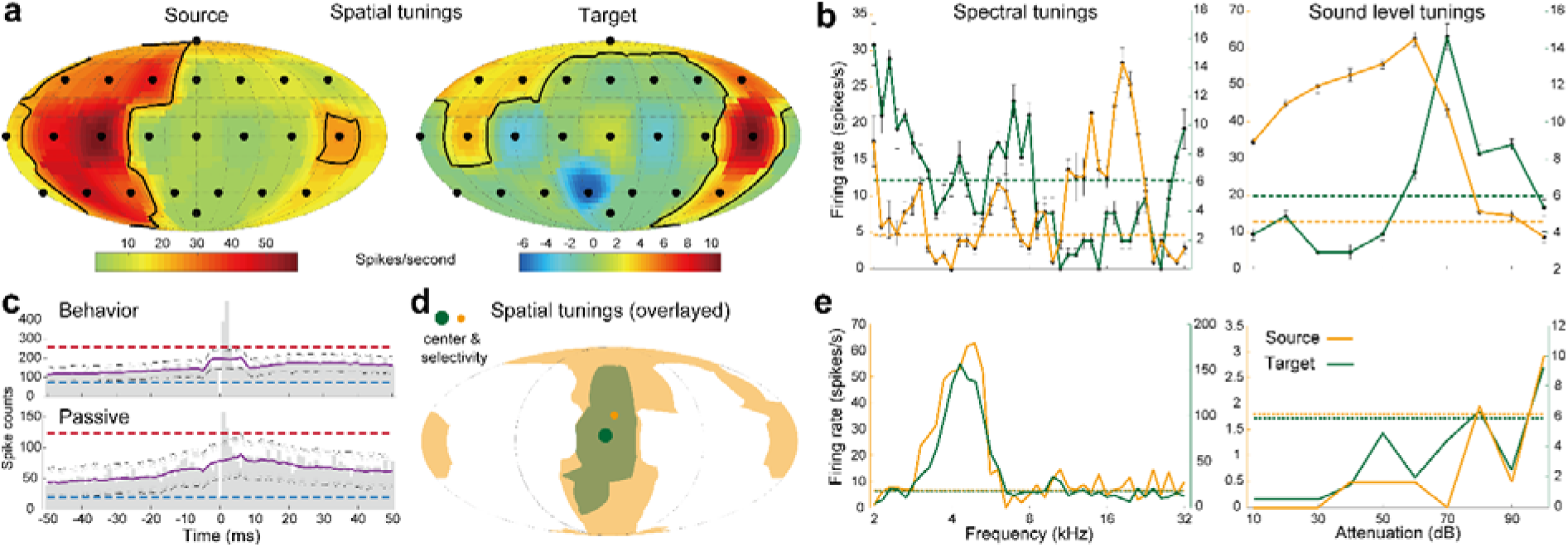
CCG is stable across brain states and consistent with sound tunings. (a) SRFs of another pair of neurons (M71V-1818). (b) Spectral and sound-level tunings of the same paired neurons shown in (a). (c) CCGs across two brain states in which the source neuron excited the target neuron. (d) Source neuron with broader spatial tuning activating a target neuron (M3T-0761) with narrower tuning. The position and size of the dots indicate the tuning center and selectivity, respectively. (e) Spectral and sound-level tunings of paired neurons (M71V-0524 and M3T-2369) with an excitatory-type CCG.

In connections with excitatory-type CCGs, we observed overlapped or matched sound tunings between source and target neurons. Similar to inhibitory-type CCGs, excitatory connections were also state independent but tended to be stronger during attention (Fig. 4c). The SRF of the target neuron was encompassed by that of the source neuron, and the two neurons had similar SRF centers (Fig. 4d). The target neuron tended to be excitatory due to its higher stimulus selectivity (dot size in Fig. 4d) and lower spontaneous firing rate (Supplementary Fig. 4a). Spectral tunings from two neurons with an excitatory connection were also highly overlapped (Fig. 4e, left), and both showed sustained firing to preferred tone frequencies (Supplementary Fig. 4b). Two putative excitatory neurons with excitatory connections (ACGs and CCG shown in Supplementary Fig. 4c) exhibited sparse firing (Supplementary Fig. 4d). Both neurons preferred lower sound levels, with overlapped tuning curves (Fig. 4e, right).

Together, these results show that functional connectivity–validated cell types can be further corroborated by comparing their auditory tuning properties.

### Three cell types exhibited distinct basic responses and auditory tuning properties

In addition to functional connectivity, three basic response properties further validated the classification. First, the response latency of NW-burst neurons was only 17 ms, much shorter than that of NW-nonburst neurons (28 ms) and about half that of BW neurons (Fig. 5a, Table 1). Second, the spontaneous firing rates of putative inhibitory NW-nonburst neurons were more than three times higher than those of excitatory NW-burst neurons and more than six times higher than those of BW neurons (Table 1). Finally, previous studies have shown that stimulus onset quenches response variability, measured with the Fano factor (FF, firing rate standard deviation divided by mean) (Churchland et al., 2010).

**Figure 5.**
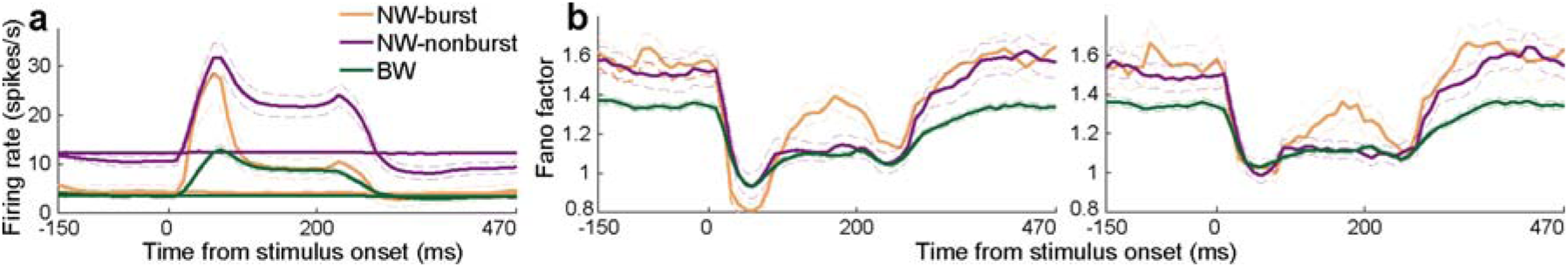
Three identified cell types exhibited distinct basic responses. (a) Raw firing rates and mean-matched firing rates (nearly flat lines) of the three cell types in response to 200 ms sound stimuli. Dashed curves indicate the standard error of the mean. (b) Fano factors of raw (left) and mean-matched (right) firing rates. Both firing rate and Fano factor were computed using a 50 ms sliding window.

**Table 1.** Summary of tuning properties for three cell types. Neurons from rostral R/RT, primary A1, and caudal CM/CL auditory cortex were combined. Analyses restricted to A1 alone yielded similar conclusions. The median recording depths were also similar across groups (500–600 µm, corresponding to upper cortical layers). First-spike latency was defined as the occurrence of spikes in three consecutive 1-ms bins.

|  | NW-burst (Ex) | NW-nonburst (In) | BW (Ex) |
| --- | --- | --- | --- |
| Num of neurons | 34 | 91 | 441 |
| Latency of first spike | **17 ms** | 28 ms | 34 ms |
| Spontaneous firing rate | 2.44 spike/s | **8.79 spike/s** | 1.39 spike/s |
| Spatial receptive field | 0.21 | **0.5** | 0.2 |
| Decoding performance | **High** | Medium | Low |
| Best location firing rate | 32 spike/s | **41 spike/s** | 19 spike/s |
| Best location (0e, 0a) | **24%** | 8% | 5% |
| Spectral bandwidth | **0.35 oct** | 0.69 oct | 0.5 oct |
| Monotonicity index | **0.23** | 0.64 | 0.54 |

Figure 5b shows the FF before (left) and after (right) matching firing rate differences. FF of spontaneous firing was similar between NW-burst and NW-nonburst neurons and higher than that of BW neurons. The larger FF in NW compared with BW neurons is consistent with previous findings in macaque V4 (Mitchell et al., 2007). During stimulus onset, NW-burst neurons showed the largest drop in FF (from 1.6 to 0.8; Fig. 5b, left) compared with the other two cell types. This difference disappeared when firing rates were matched, due to the large onset responses of NW-burst neurons. Furthermore, FF of NW-burst neurons increased again during the sustained period of stimulation, suggesting that NW-burst neurons are unreliable in encoding ongoing stimuli.

In summary, these strong response differences among the three types of neurons further validated the cell-type classification results.

Excitatory and inhibitory neurons validated through functional connectivity showed distinct sound tuning properties (Fig. 3, 4). In the final part, we compared the three cell types on their sound encoding and decoding properties. First, the sizes of SRFs in NW-burst and BW neurons were similar but only 40% of those in NW-nonburst neurons (Fig. 6a). The widths of spectral tuning were also different across cell types (Table 1). NW-burst neurons showed the sharpest tuning, with widths only half that of NW-nonburst neurons.

**Figure 6.**
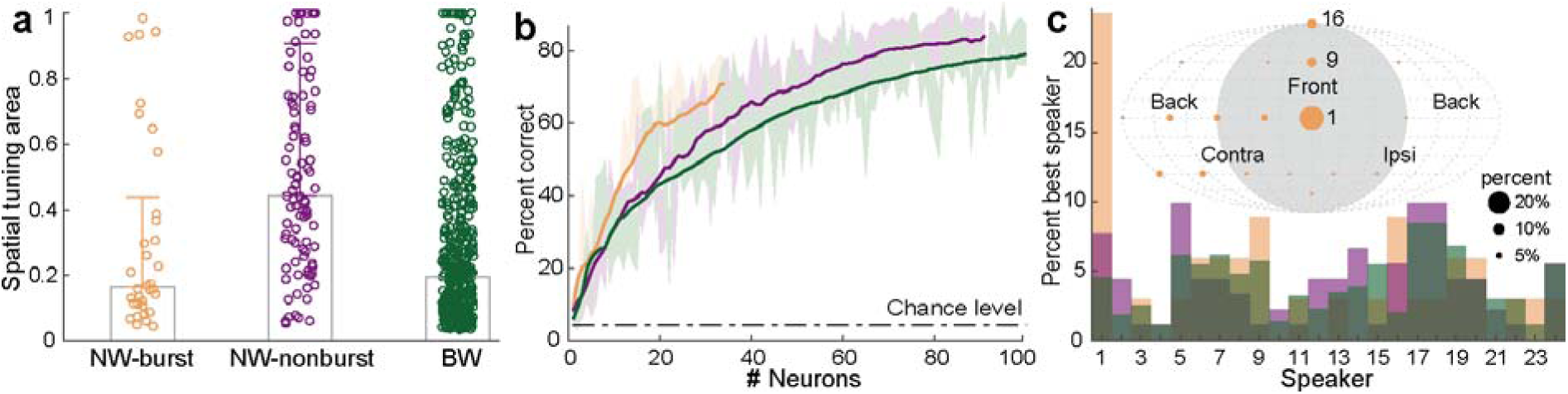
Three identified cell types exhibited distinct tuning properties to sound stimuli. (a) Areas of spatial receptive fields of individual neurons (N = 34, 91, and 441) from the three cell types. Heights of bars: median values. Error bars: standard deviations. (b) Performance of population pattern decoders using neurons from the three cell types. (c) Distribution of best speaker locations in the three cell types (N = 34, 91, and 441). Inset: overlaid best speaker locations, with dot diameter proportional to the number of NW-burst neurons preferring each speaker.

In addition to sound encoding, we compared sound location decoding performance across the three cell types (Fig. 6b). NW-burst neurons exhibited the highest decoding accuracy, while BW neurons showed the lowest. Notably, decoding performance is not necessarily positively correlated with tuning selectivity (Chen and Song, 2024); the distribution of tuning preferences also plays an important role (Belliveau et al., 2014; Chen and Song, 2024). Previous studies have compared decoding accuracy across cortical areas (e.g., rostral, primary, and caudal; Miller and Recanzone, 2009), across species such as gerbil and barn owl (Lesica et al., 2010), between auditory cortex and midbrain (Belliveau et al., 2014), across stimulus types (e.g., filtered vs. broadband noise; Wood et al., 2019), and between active and passive states (van der Heijden et al., 2018). To our knowledge, this is the first study to compare sound decoding performance among different cell types in the auditory cortex.

We found that NW-burst neurons were much more likely to prefer midline sound locations (speakers #1, 9, and 16; Fig. 6c). Specifically, 24% of NW-burst neurons had their best speaker at the direct front of the animal (0° azimuth, 0° elevation), compared with fewer than 10% in the other two cell types. Two studies in cats reported specialized auditory cortical regions with higher proportions of neurons preferring frontal sound locations than other regions (Las et al., 2008; Lee and Middlebrooks, 2013; see Discussion). However, we are not aware of any prior studies reporting differences in spatial preference among cortical cell types.

Finally, we examined sound-level tuning across the three cell types. We used the monotonicity index (MI) to quantify level selectivity, defined as the ratio of the firing rate at the highest sound level to the firing rate at the best sound level. An MI of 1 indicates that a neuron is maximally driven by the highest sound level and is highly monotonic. Nonmonotonic intensity tuning is associated with “O-shaped” frequency response areas (FRAs), which are more prominent in the auditory cortex of awake than anesthetized marmosets (Sadagopan and Wang, 2008). Our previous work showed that putative inhibitory neurons have higher MI values than excitatory neurons (Gao and Wang, 2018). Here, we observed similar results: NW-nonburst neurons and BW neurons had higher MI values (Table 1). Interestingly, NW-burst neurons had MI values only 43% of those of BW neurons, suggesting that NW-burst neurons are highly selective for sound levels.

Together, the three classified cell types were distinguished by their distinct and complementary sound-coding properties. Among them, NW-burst neurons exhibited the highest selectivity for sound location, frequency, and level.

One influential theory of auditory spatial and non-spatial processing is the “what” and “where” pathways (Romanski et al., 1999; Lomber and Malhotra, 2008; see review Rauschecker and Scott, 2009). Typically, neurons with higher spatial selectivity show lower non-spatial selectivity, and vice versa (Tian et al., 2001). However, two example NW-burst neurons exhibited the opposite trend, showing either broad or narrow tuning to both features (Fig. 7a-c). Across the population, excitatory neurons (NW-burst and BW) but not inhibitory neurons showed a significant positive correlation between spatial and spectral selectivity (Fig. 7d). Among excitatory neurons, NW-burst neurons exhibited a much stronger correlation than BW neurons (0.9 vs. 0.33). This observation is consistent with studies showing distributed sensitivity to vocalizations (“what”) in macaque auditory cortex (Recanzone, 2008) and modulation of responses by both spatial and non-spatial attributes in macaque prefrontal cortex (Cohen et al., 2004).

**Figure 7.**
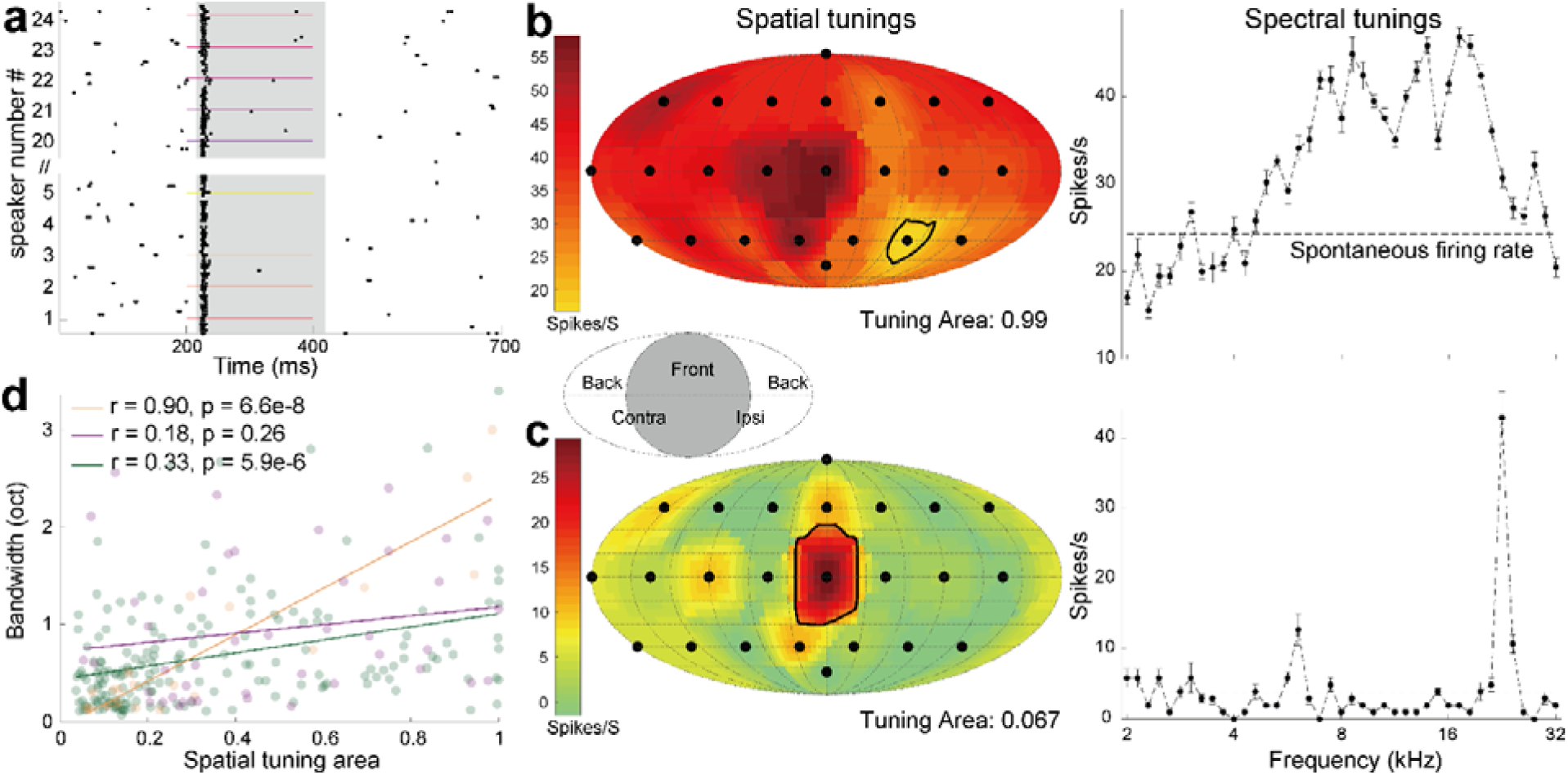
Correlated spatial and spectral selectivity. (a) Spike raster of an example NW-burst neuron (M3T-2412) that was unselective to speaker locations and fired bursts of spikes at the stimulus onsets. (b) Example neuron in (a) was broadly tuned to both sound location and frequency. (c) An NW-burst neuron (M3T-0902) that was highly selective for both features. (d) Scatter plot of receptive field size for sound frequency (Y-axis) versus location (X-axis) (N = 20, 43, and 184). Thick lines represent linear regression fits.

Because pure tone frequency is simpler than vocalizations, the “what” selectivity metric used here may be suboptimal. To address this, we further tested 20 marmoset vocalization tokens in 103 units (Fig. 8). Due to the small sample size, we did not distinguish among cell types. Using the monkey call index, as in previous studies (Tian et al., 2001; Recanzone, 2008), we quantified vocalization selectivity. Consistent with frequency selectivity, vocalization selectivity was also significantly correlated with spatial selectivity (p < 1e-7). Moreover, the explained variance (r² = 0.495) was even larger than that for frequency selectivity (r² = 0.33, BW neurons, the majority).

**Figure 8.**
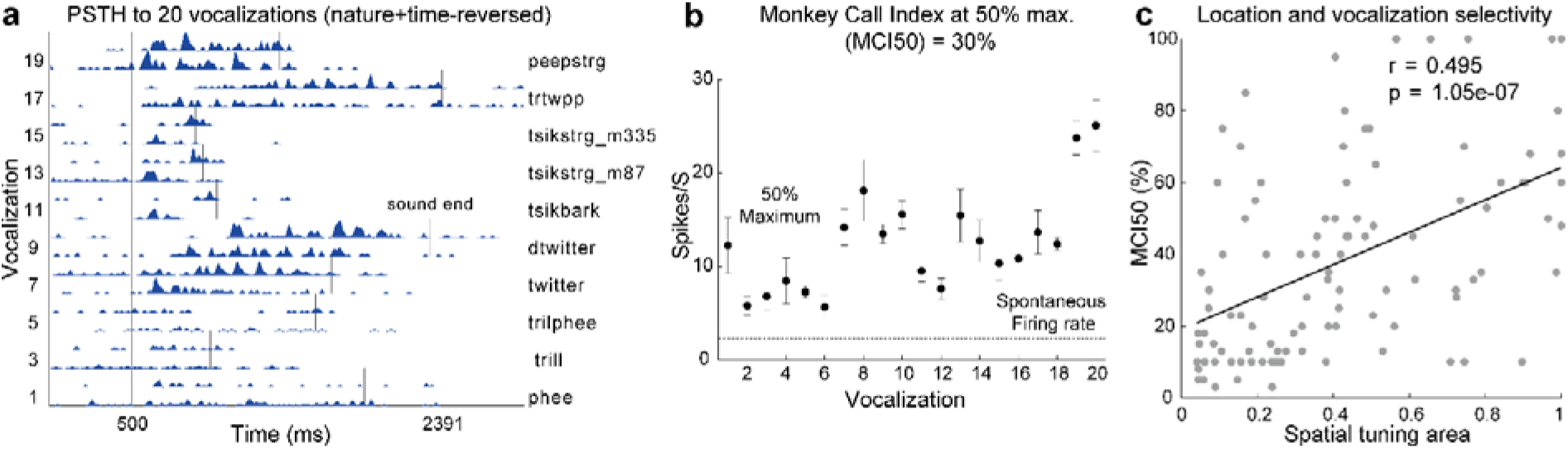
Correlated spatial and vocalization selectivity. (a) PSTH of an example neuron (M6X-0877) to 20 vocalizations. There were 10 types of vocalizations, and each one was presented either in the normal or time-reversed order. PSTH during each vocalization was normalized to have a peak value of 1. All vocalizations were aligned at the beginning (500 ms), and each one has a different duration. (b) The averaged firing rate of five trials at 20 vocalizations of same unit. The unit is significantly tuned to the vocalizations (p-value is 1.3e-10, ANOVA). Firing rate from six vocalizations was higher than the 50% maximum threshold (red dashed line), so the monkey call index at 50% maximum (MCI50) was 30% (Tian et al., 2001; Recanzone, 2008). The error bar indicates the standard error of the mean. (c) Scatter plot of normalized spatial tuning area (X-axis) against MCI50 for all the 103 units tested with both types of stimuli.

Together, these observations of positively correlated spatial and non-spatial selectivity raise further questions about the auditory dual-stream hypothesis (see reviews by Bizley and Walker, 2009; Recanzone and Cohen, 2010).

## Discussion

In this study, we analyzed over 1800 single units recorded from auditory cortex of passively or actively listening marmosets and classified cell types based on spike waveform width and firing patterns. We defined three cell types: NW-burst, NW-nonburst, and BW. We validated these classifications by showing that NW-burst and BW neurons were excitatory, while NW-nonburst neurons were inhibitory. One independent method supported this conclusion: functional connectivity between paired neurons. Finally, we compared sound-tuning properties among the three cell types and found that excitatory NW-burst neurons exhibited the fastest and most selective responses to sound stimuli. Below, we discuss three major aspects of our findings: (1) the classification methods, (2) the validation methods, and (3) the auditory tuning properties.

### Methods for classifying cell types in primates

Primate cell-type classification based on neurophysiological recordings can be broadly categorized into three approaches: (1) spike waveform features, (2) spike firing patterns, and (3) unsupervised data-driven clustering methods.

Waveform-based classification. Putative inhibitory and excitatory neurons are most commonly distinguished by the width of their spike waveforms, with NW corresponding to inhibitory neurons and BW corresponding to excitatory neurons, based on trough-to-peak duration. In some studies, the boundary between NW and BW neurons is clear (Onorato et al., 2020; Tsunada et al., 2012), while in others it is less distinct (Gao and Wang, 2018; Trainito et al., 2019; Selvanayagam et al., 2024). The reported proportion of NW neurons also varies widely across studies: 28% of neurons in macaque V4 (Mitchell et al., 2007), 71% in macaque V1 (Onorato et al., 2020), and 47% in nonprimary auditory cortical areas (Tsunada et al., 2012). In addition to trough-to-peak duration, other waveform features have been used, including half-amplitude duration (Barthó et al., 2004; Lee et al., 2021), spike peak ratios (Lee et al., 2021), repolarization slope (Jia et al., 2019), and repolarization time (Ardid et al., 2015). Despite these alternatives, trough-to-peak duration remains the most widely applied criterion for distinguishing NW and BW neurons in primate studies.

In addition to spike waveform, neuron’s firing statistics, especially burst firing, have also been widely used as markers of cell type. In the cortex, two types of bursting neurons have been described. Intrinsic bursting (IB) neurons are found in both superficial (de Kock and Sakmann, 2008; Senzai et al., 2019; Wang et al., 2020) and deep cortical layers (Williamson and Polley, 2019) across many species. In contrast, “chattering” or NW-burst neurons are primarily located in the superficial layers (though see Cardin et al., 2005) of larger animals such as cats (Gray and McCormick, 1996; Nowak et al., 2003) and monkeys (Onorato et al., 2020; Liu and Wang, 2022). IB neurons typically fire dense spike trains and are broadly tuned to sensory features (Sakata and Harris, 2009; Sun et al., 2013). By contrast, NW-burst neurons in macaque superficial layers exhibit much higher selectivity for sensory features (Onorato et al., 2020). In this study, we separated NW neurons into NW-nonburst and NW-burst subtypes and found that they exhibit substantially different tuning properties. This suggests that bursting is a reliable and functionally meaningful marker for distinguishing cortical cell types.

The last approach is the unsupervised data-driven method. There are two main strategies. The first still relies on waveform features (at least two), such as trough-to-peak duration and repolarization potential, but defines cell-type boundaries using Gaussian mixture models. Using this approach, Trainito et al. (2019) identified four neuronal clusters across three cortical regions of the macaque and demonstrated distinct functions among them. Similarly, Liu and Wang (2022) identified three clusters of neurons with this method, and the clustering results were highly consistent with feature-based classifications. The second strategy uses nonlinear dimensionality reduction to extract features from the entire waveform rather than human-defined metrics. The most widely applied techniques are t-distributed stochastic neighbor embedding (t-SNE) and uniform manifold approximation and projection (UMAP). Jia et al. (2019) applied t-SNE in the mouse visual cortex and revealed three neuronal clusters. Using optogenetic tagging, they confirmed that all 29 PV neurons belonged to the NW cluster. In another example, Lee et al. (2021) applied UMAP in the macaque premotor cortex and identified eight clusters. Recently, deep learning methods have been introduced for cell type clustering (Vishnubhotla et al., 2023; Chen et al., 2024; Beau et al., 2025; Yu et al., 2025). In this study, we applied unsupervised K-means clustering to set the boundary between burst and nonburst neurons (Supplementary Fig. 1d).

### Methods for validating cell type classification

Cell types classified through neurophysiological recordings can be validated against ground truth using several approaches:

Morphological reconstruction. One method is in vitro reconstruction of recorded neurons following juxtacellular recordings (de Kock et al., 2007; Sakata and Harris, 2009). For example, Gray and McCormick (1996) demonstrated that “chattering” neurons in cat V1 exhibited pyramidal-like morphology. However, this approach requires sacrificing the animal after recording a single neuron, and thus is rarely applied in primate studies.

Antidromic stimulation. Another method is antidromic stimulation of the axon terminals of excitatory projection neurons, using either electrodes (Johnston et al., 2009; El-Shamayleh et al., 2013) or light (Williamson and Polley, 2019). For example, NW neurons in macaque motor cortex were shown to be excitatory using antidromic tagging (Vigneswaran et al., 2011). The limitation of this method is its low yield, and it cannot identify neurons that lack long-range projections.

Functional connectivity. A third method is functional connectivity analysis based on excitatory–inhibitory interactions measured with CCG (Constantinidis et al., 2002; Peyrache et al., 2012; Jia et al., 2013). However, this approach requires a large number of postsynaptic spikes to achieve statistical significance (Gerstein and Perkel, 1972). It also demands long recording sessions or neurons with high firing rates. In addition, CCG-based connections may not always reflect monosynaptic inputs and can be confounded by common drive (English et al., 2017).

Optogenetics tagging. A fourth method is optogenetics tagging of genetically defined cell types, typically performed in transgenic mice (Keller et al., 2018; Lakunina et al., 2020; Onorato et al., 2025). Extension of this method to non-rodent species requires viral vectors carrying cell type–specific promoters for excitatory (Nurminen et al., 2018; Jendritza et al., 2023) or inhibitory (De et al., 2020; Town et al., 2023) neurons. A major limitation is that expression of cell type–specific markers are technically challenging in nonhuman primates. Moreover, untagged neurons may represent false negatives because opsins may fail to label all target cell types, and light penetration is limited in middle and deep cortical layers (Town et al., 2023).

In this study, we validated cell types classification using functional connectivity. Importantly, we corroborated these validations by comparing receptive fields between paired neurons and showing that their similarities or complementarities were consistent with excitatory or inhibitory connections, respectively.

### Cell type specific spatial and non-spatial tuning properties

How different regions of the auditory cortex contribute to spatial processing has been studied extensively. In primates, caudal areas are more sharply tuned for sound locations than primary (A1) and rostral areas (Tian et al., 2001; Woods et al., 2006; Zhou and Wang, 2012; Remington and Wang, 2019). In cats, 28% of space-selective neurons in the posterior anterior ectosylvian sulcus (AES) had strongly modulated frontal receptive fields (–30° to 30°), compared with 8% in A1 and 17% in anterior AES (Las et al., 2008). Among neurons with measurable centroids, 58.5% in the dorsal zone (DZ) were located in the frontal quadrant (–45° to 45°), compared with only 35.8% in the posterior auditory field (PAF) (Lee and Middlebrooks, 2013). In contrast, much less is known about the contributions of different cell types to spatial processing. Only two relevant rodent studies exist. One study reported that inhibitory neurons in the mouse inferior colliculus are sharply tuned to sound localization cues (Chen and Song, 2024). Notably, inhibitory neurons in the IC are also sharply tuned to sound frequency (Chen and Song, 2019) and exhibit spike waveforms similar to excitatory neurons (One et al., 2017). Another study found that putative inhibitory neurons in the gerbil auditory cortex are broadly tuned to sound locations (Amaro et al., 2022). NW-nonburst neurons (inhibitory) showed broad tuning, while BW neurons (excitatory) showed narrow tuning, are consistent with previous results in auditory cortex (Amaro et al., 2022).

The tunings of NW-burst neurons are also consistent with two previous studies. NW-burst neurons in macaque V1 are more selective for grating orientation and their responses are phase-locked to drifting grating stimuli (Onorato et al., 2020). Similarly, high-frequency bursting neurons (their Bu1 and Bu2) in marmoset auditory cortex are also phase-locked to time-varying acoustic signals (Liu and Wang, 2022). These two findings are similar to what we observed in NW-burst neurons: higher auditory spatial/spectral tunings and stronger onset/offset responses. Why do NW-burst neurons exhibit sharp and correlated spatial-spectral tuning? We suspect they receive direct inputs from bottom-up auditory thalamus. One piece of evidence is their much shorter response latency compared to the other two types of neurons. Another piece of evidence is that neurons in auditory thalamus have narrower spectral tuning than cortical neurons (Bartlett et al., 2011). It is unlikely that this sharp tuning arises from local inhibitory cortical circuits. Lateral inhibition is mainly observed during sustained rather than onset responses (Sadagopan and Wang, 2010; Bartlett et al., 2011), and likely contributes to the longer latency (34 vs. 17 ms) and sustained responses observed in BW neurons (Fig. 5a). In contrast, NW-burst neurons primarily fire at the onset of sound stimuli (Liu and Wang, 2022).

Together, NW-burst neurons may use phase-locked spike timing to encode temporally modulated stimuli and firing rate to encode sound location, frequency and intensity with high selectivity and short latency. Given these properties, NW-burst neurons should play a pivotal role in coding and transmitting stimulus information.

## Acknowledgements

Support was contributed by US National Institutes of Health grants DC003180 (X.W.).

## Supplementary Figures

**Supplementary Figure 1.**
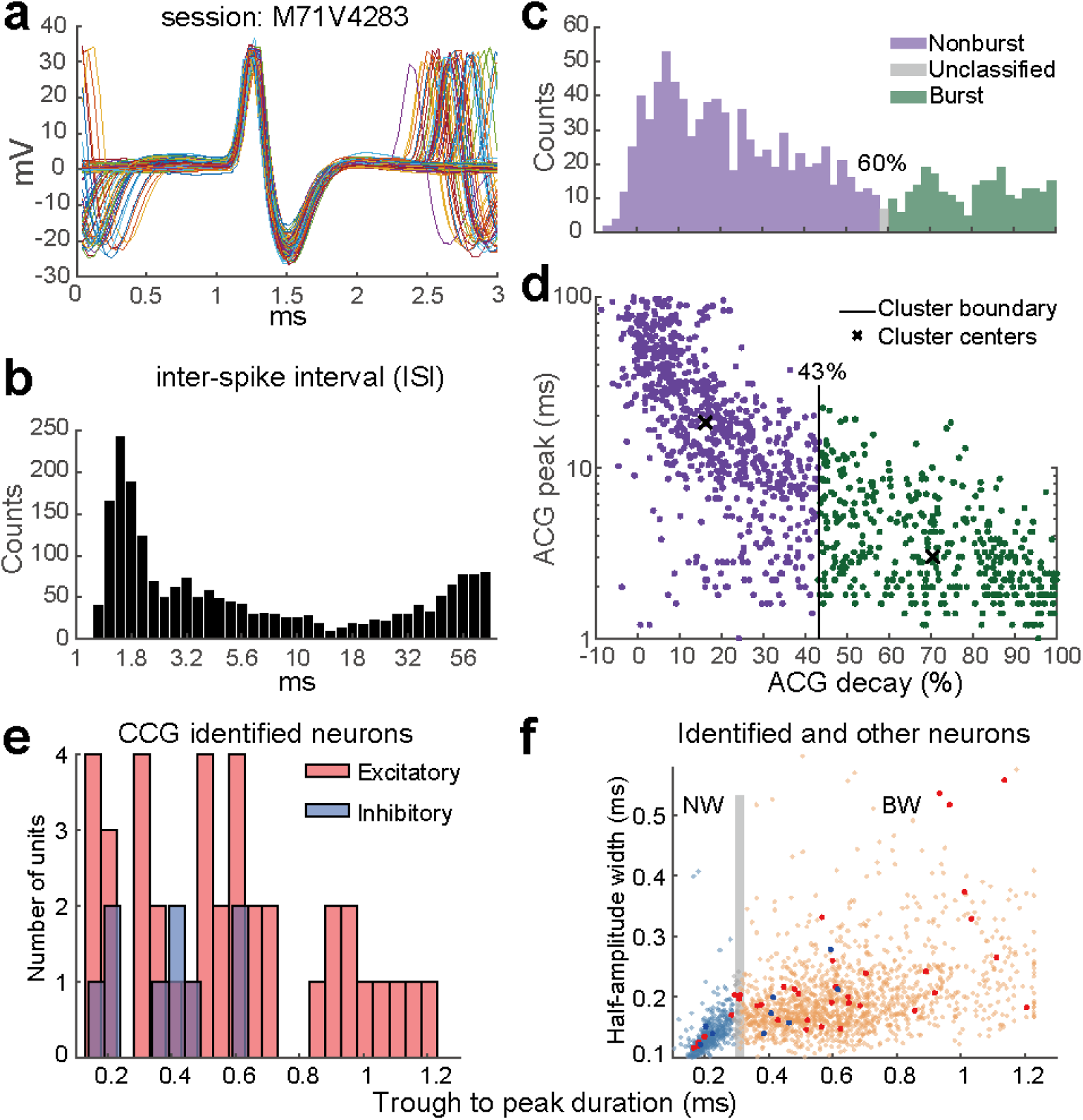
Classification of nonburst and burst neurons (related to Fig. 1). (a) There were 100 spike waveforms from one NW-burst neuron that were aligned by the trough of the waveform at 1.5 ms. Since this neuron fired burst spikes within a short period, the spike waveforms before and after the center-aligned spike were also captured. (b) The histogram of an example ISI with a sharp peak at 1.7 ms (4637 spikes in total). Notice the X-axis is log10. (c) The histogram of ACG decay for nonburst (purple) and burst (green) neurons. (d) The scatter of ACG decay (X-axis) versus ACG peak (Y-axis). (e) Histogram of spike width (trough to peak duration) for identified excitatory and inhibitory neurons using CCG in marmosets. (f) Scatter plot of two metrics of spike width (trough to peak and half amplitude) in NW and BW neurons (half-transparent cyan and orange) with overlayed inhibitory and excitatory neurons (blue and red). The X-axis is the same as (e). Six out of the nine identified inhibitory neurons have a BW (66.7%), and 29 out of the 37 excitatory neurons have a BW.

**Supplementary Figure 2.**
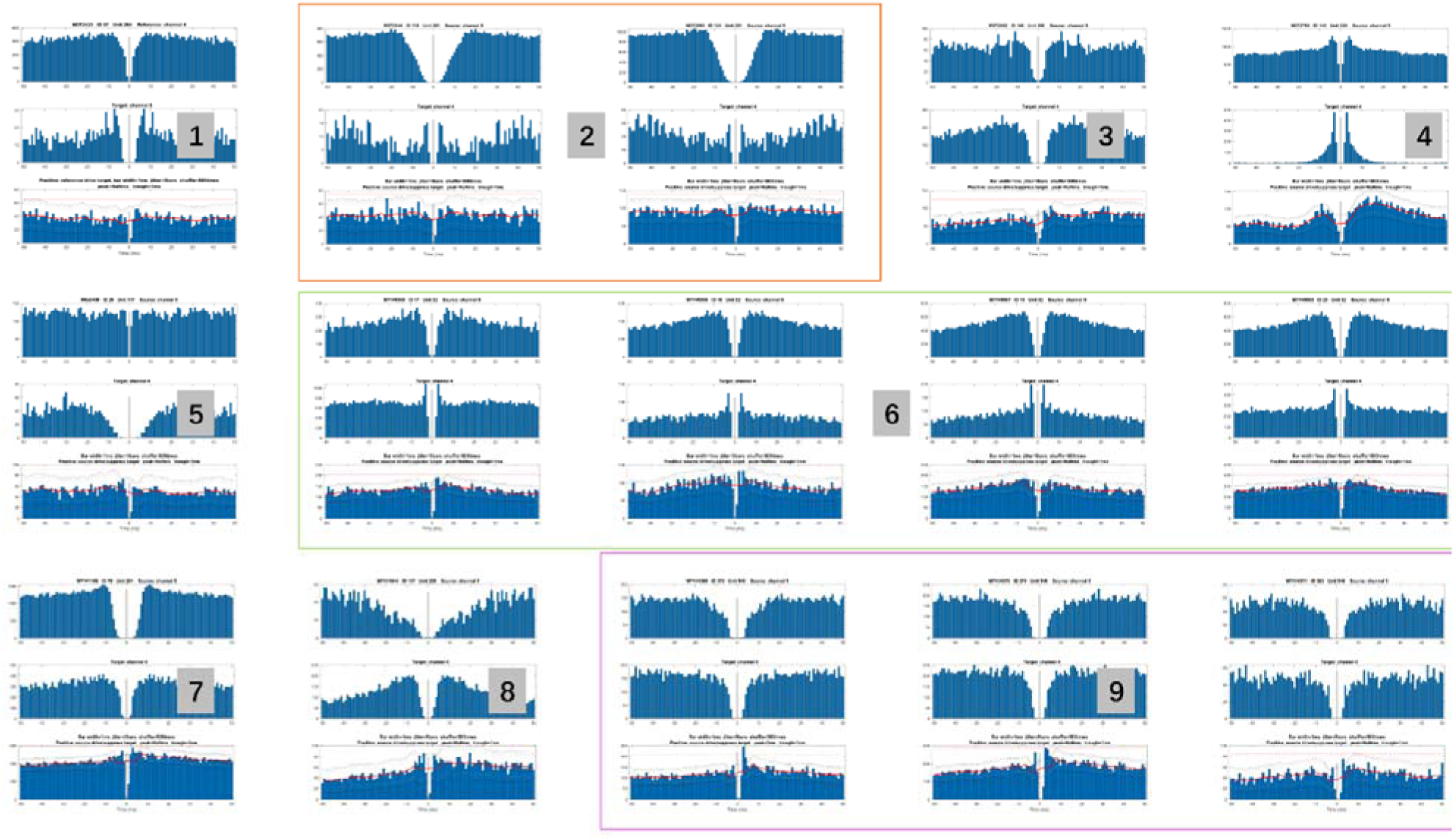
Auto- and cross-correlogram (ACG and CCG) of nine identified inhibitory neurons (related to Fig. 2a, b). Each neuron can have over one session (e.g., four sessions in #6). The top and middle rows show the ACG of presynaptic and postsynaptic neurons, respectively. The bottom row shows the CCG between connected neurons.

**Supplementary Figure 3.**
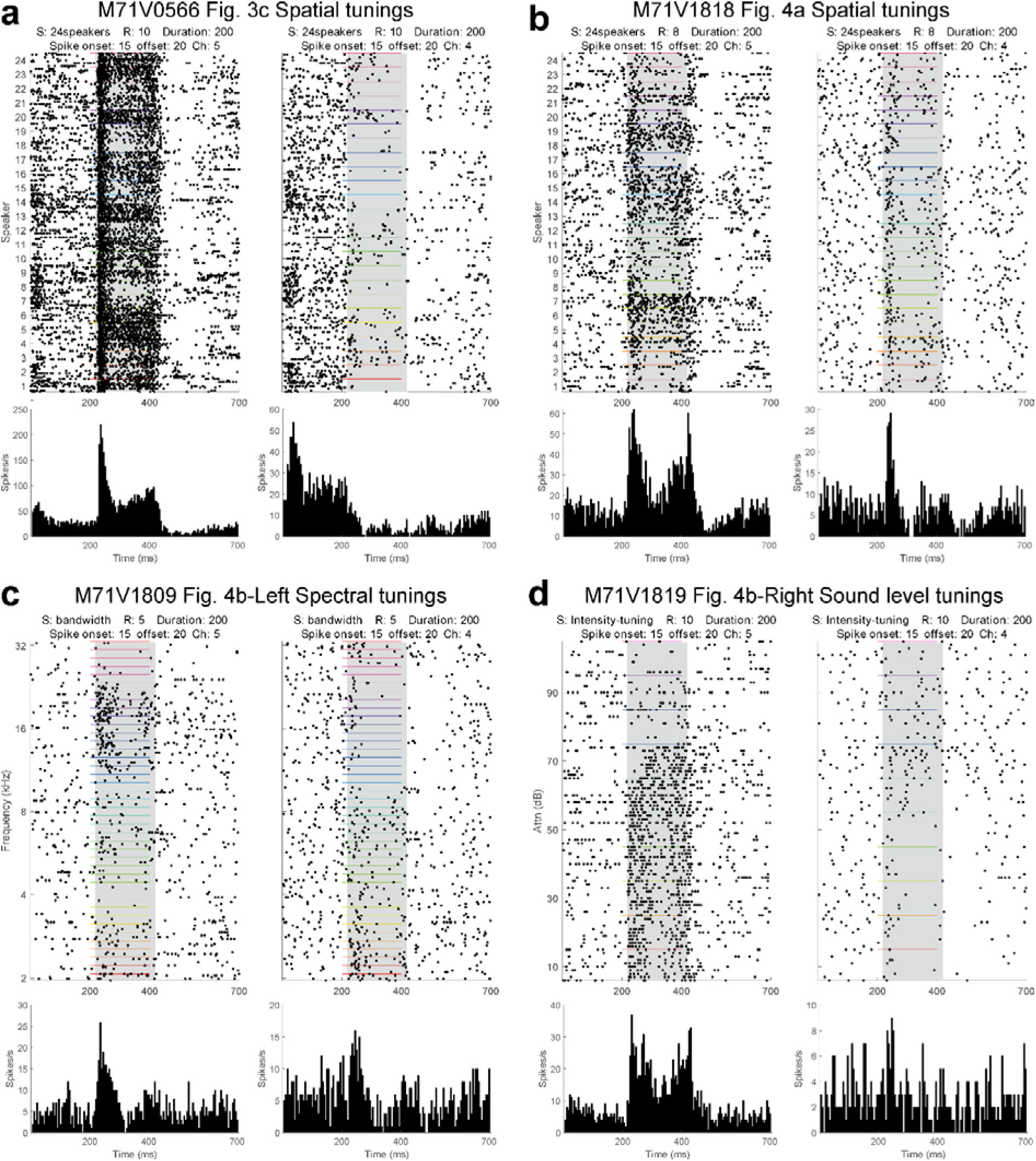
Spike raster and PSTH in paired neurons from four sessions (related to Fig. 3c, 4a, b). (a-d) Spike rater and PSTH of four example sessions shown in Fig. 3c, 4a, 4b-left, and 4b-right.

**Supplementary Figure 4.**
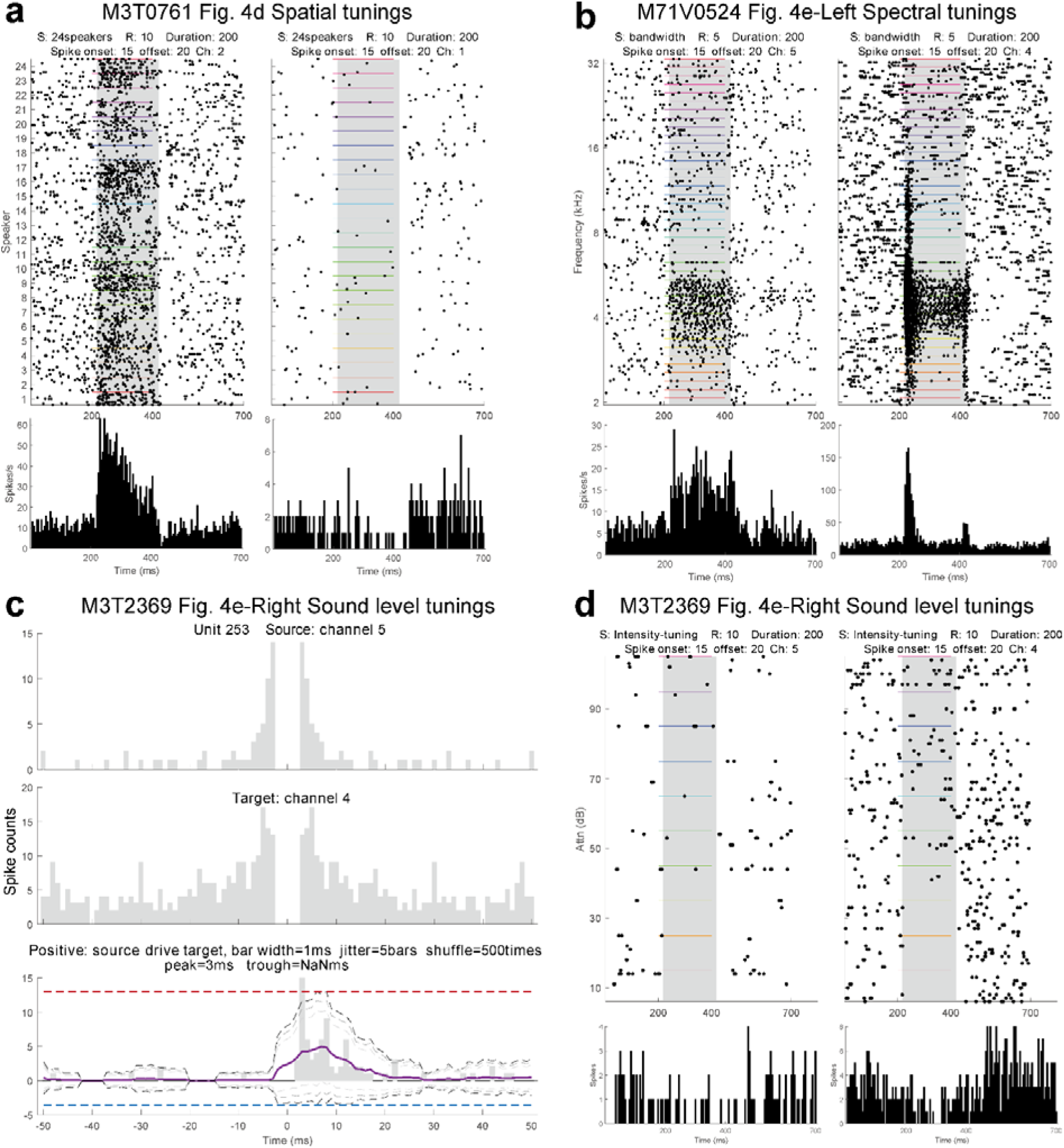
Spike raster, PSTH, ACG/CCG in paired neurons from three sessions (related to Fig. 4d, e). (a, b, d) Spike rater and PSTH of three example paired neurons shown in Fig. 4d, 4e-left, and 4e-right. (c) ACG and CCG for the neuron pair in (d).

